# Housekeeping genes are enriched in rheumatoid arthritis-related genes

**DOI:** 10.1101/2025.02.22.639692

**Authors:** Wenhua Lv, Hao Tang, Guanglong Feng, Shuhao Zhang, Zhenwei Shang, Mingming Zhang, Rui Chen, Xiangshu Cheng, Xin Meng, Hongchao Lv, Ruijie Zhang

## Abstract

Housekeeping genes are often considered to be stably expressed in any tissue, cell, or disease state, and the relationship between housekeeping genes and rheumatoid arthritis (RA) has never been systematically analyzed. Therefore we aimed to explore whether housekeeping genes are abnormally expressed in RA by analyzing 15 datasets that were filtered from 140 gene expression profiles. By cumulative hypergeometric test, for 73.3% (11/15) of the datasets, housekeeping genes were significantly (p < 0.05) enriched in differentially expressed genes (DEGs) between RA patients and healthy controls. Using the relative enrichment factor (Re) as an effect size, a Meta-analysis strategy was adopted to integrate the individual results. The logarithmic Re estimation for multiple studies was 0.48 with the 95% confidence interval ranging from 0.17 to 0.80. The combined effect size was significantly higher than that of the randomized case (p = 0.0012) suggesting a consistent result with the hypergeometric test. The robustness of results was verified by sensitivity and subgroup analyses. Furthermore, the expression fluctuation of housekeeping genes was higher in RA patients than healthy controls and the correlation between DNA methylation and gene expression of housekeeping genes was significantly higher than that of non-housekeeping genes, which indicates the potential epigenetic regulation relationship between housekeeping genes and the pathogenesis of RA. This study identified for the first time a significant enrichment of housekeeping genes in DEGs of RA, which might provide new perspectives for understanding the potential role of housekeeping genes in RA.

## Introduction

Rheumatoid arthritis (RA) is a chronic destructive inflammatory autoimmune disease, with broader clinical complications, including comorbidities that particularly affect bone, vasculature, and metabolic function and cognition throughout the body[1]. In recent years, there have been many large scale studies focusing on the molecular level of RA, including genome wide association studies (GWAS)[2, 3], gene expression regulation studies[4] and epigenome wide association studies (EWAS)[5, 6]. Several risk SNPs, CpG sites, risk genes and risk pathways associated with RA were identified, which made important contributions to the elucidation of the molecular mechanism of RA. However, it is not possible to clearly distinguish RA patients from healthy controls or clearly divide RA patients into different subtypes. In addition, patients with RA respond differently to medication therapy[7], suggesting that RA is a chronic disease with a very complex mechanism. As one of the complex diseases, RA is characterized by genetic micropotency, that is, the common dysregulation of hundreds of genes leads to the occurrence of diseases, while the contribution of each gene is very weak[8]. Therefore, we hypothesize that RA may be caused by abnormalities in various genes that play essential roles in maintaining the body’s normal functioning.

Housekeeping (HK) genes are genes that are needed to maintain the basic cellular functions that are essential for the survival of a cell, regardless of its specific role in the organism or tissue. Thus, human HK genes are supposed to be expressed in all cells under normal conditions, irrespective of cell cycle state, tissue type and developmental stage[9, 10]. Due to the relative stability of expression levels across tissues and organs, HK genes were often used as controls to quantify the expression levels of other genes[11, 12]. In recent years, there have been researches that observed significant relationship between HK genes and complex diseases. For instance, ATP Binding Cassette Subfamily F Member 1 (ABCF1) is upregulated in hepatocellular carcinoma[13], expression levels of AdipoR1 and AdipoR2 are significantly elevated in gastric cancer[14]. Over-expression of MAGT1 is associated with invasiveness and poor prognosis of colorectal cancer[15]. The abnormal expression of HMGB1 is thought to be related to the pathogenesis of autoimmune diseases such as systemic lupus erythematosus and rheumatoid arthritis[16]. However, so far there have been no large-scale studies systematically exploring the potential alteration of HK genes during the process of complex diseases.

Hence we are interested in the role of housekeeping genes in rheumatoid arthritis. We collected all available transcriptome data of both RA and healthy control individuals in public databases and identified differentially expressed genes (DEGs) for each dataset. Next we intended to investigate the relationship between HK genes and DEGs through enrichment analysis. Then the classic Meta-analysis method was used to integrate and validate the observations in this research. Our study explored transcriptional changes of HK genes in RA for the first time and may provide new insights into further interpretation of RA.

## Materials and Methods

### Acquisition of transcriptome data of RA

Transcriptome data of RA were obtained from the Rheumatoid Arthritis Bioinformatics/Big data Center (RABC) which is the first large data resource platform providing data storage, processing, and analysis for RA research[17]. To ensure the reliability of the analysis results, only transcriptome data with a sample size greater than 20 were included. To avoid the effect of drug therapy and other treatments on the patient’s gene expression, only baseline RA individuals prior to treatment were included. The criteria and process of data inclusion and exclusion are shown in Figure 1. Finally, 15 sets of transcriptome data including both RA and healthy control were included in this study. The level 3 expression matrix data were downloaded from RABC. The clinical information for both RA patients and healthy control individuals are provided in Table S1. **HK gene set**

**Figure 1.**
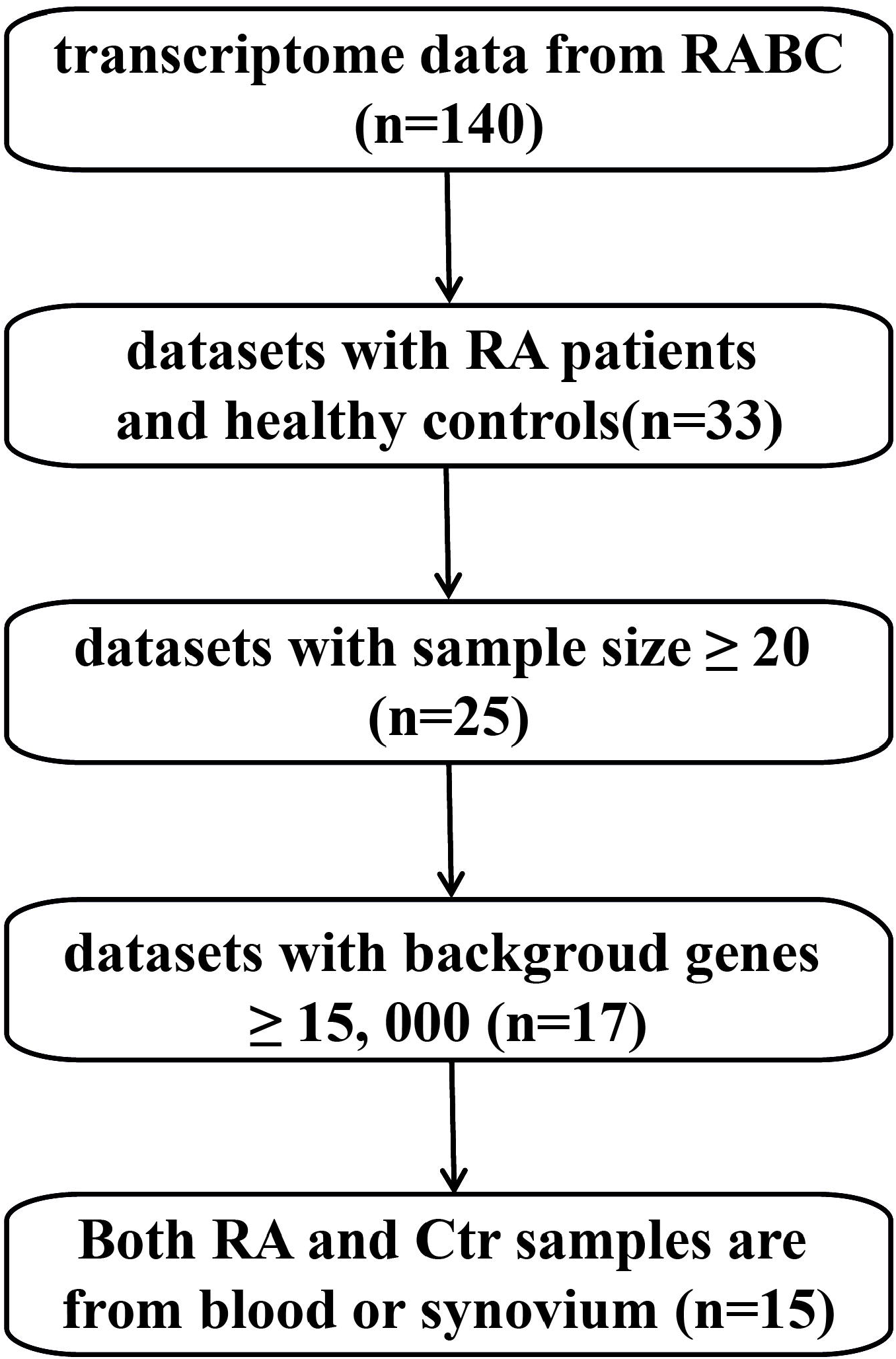
The flow chart of dataset inclusion and exclusion.

The whole human HK gene list with high confidence including 3804 HK genes was obtained from a classic study that identified HK genes based on high-throughput sequencing data[18]. They set three criteria to screen exons with uniform expression level across different tissues : (i) expression was observed in all tissues; (ii)low variance over tissues; and (iii) no exceptional expression in any single tissue. A gene was defined as housekeeping gene if at least one RefSeq transcript has more than half of its exons meeting the previous criteria.

### Differential expression analysis between RA and healthy control

Limma package[19] in R was utilized to perform differential expression analysis. For different transcriptome datasets, we set a common criteria to identify the differentially expressed genes (DEGs) between RA and healthy control group. A gene is defined as a DEG if it ranks at the top 20% of the whole gene list sorted by absolute logarithmic fold change (|logFC|) in descending order and the p-value of the t-test is less than 0.05.

### Enrichment analysis between HK gene sets and DEG sets

For each independent RA dataset, the number of DEGs n_g_, the number of HK genes annotated to the background gene set n_G_, the number of overlap between DEGs and HK genes n_k_, and the number of background genes n_N_ were calculated. The enrichment p value was defined by the cumulative hypergeometric formula (Formula 1) inflecting the probability that the size of interactions between HK gene set and n_g_ genes randomly extracted from the whole background genes is greater than n_k_.

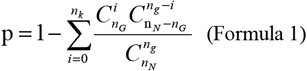

### Calculation of effect size of enrichment degree between DEGs and HK genes

For clarity, DEGs and HK genes are denoted by the letters A and B, respectively. According to the principle of independence, the probability of product events (AB) is equal to the product of the probability of A and the probability of B. Specifically 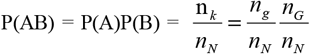. We used relative enrichment factors (Re) [20] to reflect the degree to which DEGs were enriched to HK gene set. The Re value is calculated according to the value in Table 1 (Formula 2). The Re value is equal to 1 if A and B are independent. Conversely, the greater the degree of positive deviation of the Re value from 1, the greater the enrichment of DEGs to the HK gene set. As we know, in all statistical computations it is best to transform x into its logarithm which avoids difficulties due to asymmetry. The log x transformation will retain its numerical value, merely changing in sign. Moreover, the sampling variance of log x is a very simple expression free of’nuisance parameters. This is especially true if one transforms into y=log_e_(x). Therefore, log_e_(Re) was treated as effect size in this study. Based on the Woolf method[21], the sampling variance of log_e_(Re) is represented by Var(log_e_(Re)) that is equal to 1/n_k_ + 1/n_N_ + 1/n_g_+ 1/n_G._ Then lower and upper limits of the 95% confidence interval for the effect size were defined according to the Formula 3 and Formula 4.

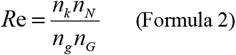

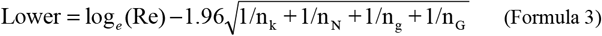

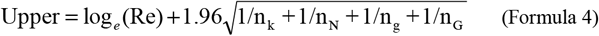

**Table 1.** Two-by-two contingency table for DEGs and HK genes.

| | DEGs (A) | Non-DEGs ($\bar{A}$) | Total |
| --- | --- | --- | --- |
| HK genes (B) | $n_k$ | $n_G - n_k$ | $n_G$ |
| Non-HK genes ($\bar{B}$) | $n_g$ | $n_N - n_g - (n_G - n_k)$ | $n_N - n_G$ |
| Total | $n_g - n_k$ | $n_N - n_g$ | $n_N$ |

### Integration of enrichment results for multiple studies based on Meta-analysis strategies

An inverse-variance weighted Meta-analysis[22] was conducted using Metafor package in R software with Re values for each individual study. Heterogeneity across different studies was evaluated by p value and I^2^ values from the heterogeneity test. A p value less than 0.05 or I^2^ value greater than 50% indicates large heterogeneity. Depending on the size of heterogeneity, effect sizes from multiple individual studies were pooled using either a fixed-effect model or a random-effect model. Pooled effect size and heterogeneity results were visualized by forest plots. The publication bias was evaluated by funnel diagram, Begg’s and Egger’s tests, where p value less than 0.05 indicates significant publication bias. The stability of Meta-analysis was assess by a sensitivity analysis which removed an individual study from the whole list each time. All datasets were classified based on the tissue source and a subgroup analysis was conducted for blood sample datasets.

To validate the reproducibility, another standard error estimation method for the effect size (Re) was applied to this research. Specifically, 100,000 random perturbations were conducted to generate hypergeometric background distribution of Re for each of the 15 datasets and then the standard errors were calculated. The 95% confidence interval for the effect size was then calculated and integration of effect size for each dataset was performed by Meta-analysis.

## Results

### An overview of the research design

A total of 140 transcriptome datasets are included in the RABC database. We screened these transcriptome datasets according to the criteria described in Figure 1 (details see Materials and methods). First, 111 datasets were removed due to the lack of healthy control individuals. Second, 8 datasets were removed due to the total sample size less than 20. Third, 8 datasets were removed since the number of background genes less than 15, 000. Fourth, 2 datasets were removed due to the different tissue orgins of RA and control samples, for instance RA samples were derived from synovium while control samples were derived from whole blood. At last, 15 transcriptome datasets were included in this research. The detailed information for each datasets that meets the criteria is shown in Table 2.

**Table 2.**
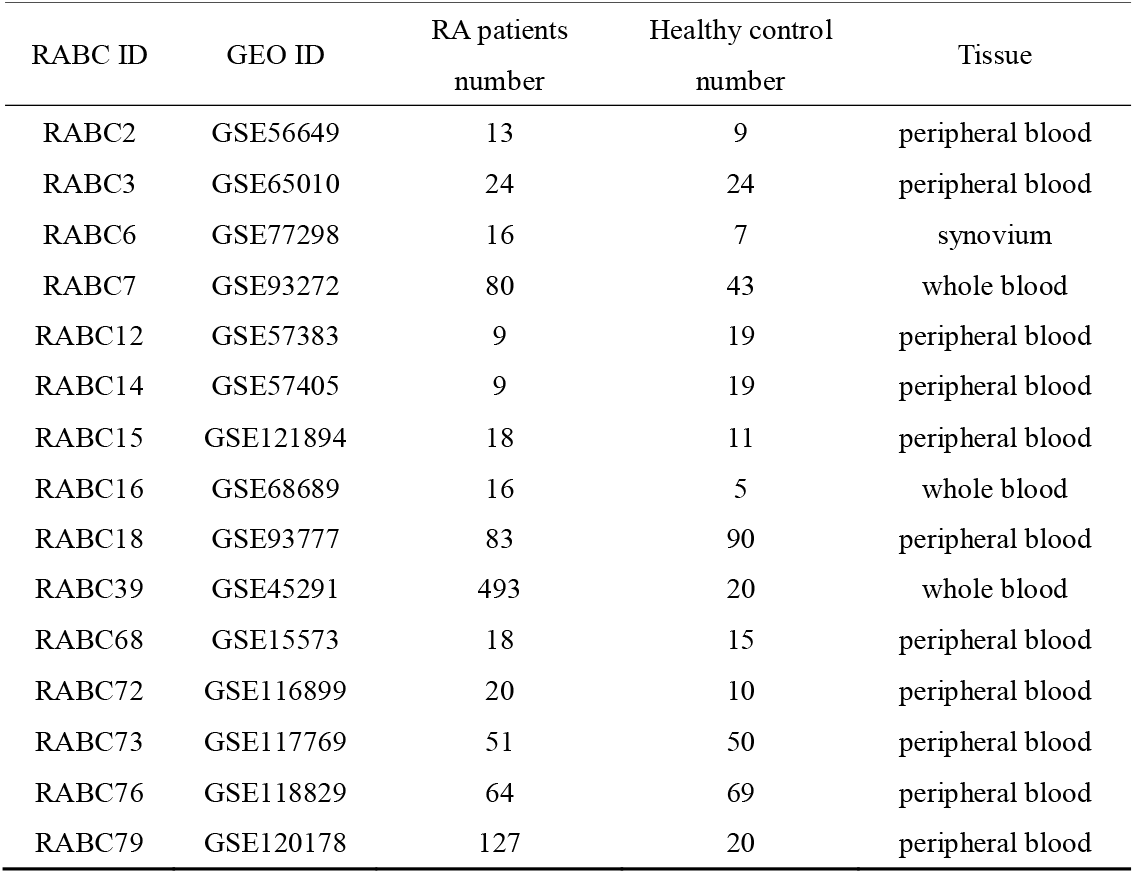
Detailed information of datasets included in this research.

### DEGs between RA and healthy control are enriched to HK genes

For each individual transcriptome datasets, DEGs between RA patients at baseline and healthy control were identified. Then the overlap between DEGs and HK genes annotated to background were calculated. For most datasets, housekeeping genes were significantly enriched into DEGs with the p-value of cumulative hypergeometry test less than 0.05. This trend is even more pronounced in studies with large sample sizes (Table 3). To reflect the degree that HK genes were enriched to DEGs, the point estimation and the 95% confidence interval of Re were calculated (see Methods and materials). Ln(Re) values ranges from −0.51 to 0.88 for these individual studies and 80% (12/15) studies have ln(Re) values greater than 0. This result further quantificationally confirms the fact that HK genes are significantly enriched in RA-related genes.

**Table 3.** Enrichment result based on cumulative hypergeometric tests.

| RABC ID | Sample<br>size | Overlap<br>gene<br>($n_k$) | HK<br>gene($n_G$) | Non-HK<br>gene<br>($n_N - n_G$) | DEGs<br>($n_g$) | P value | $\ln(\text{Re})$ | Lower<br>limit | Upper<br>limit |
| --- | --- | --- | --- | --- | --- | --- | --- | --- | --- |
| RABC2 | 22 | 607 | 3610 | 18045 | 1995 | 0 | 0.60 | 0.50 | 0.70 |
| RABC3 | 48 | 159 | 3610 | 18045 | 768 | 1.69E-03 | 0.22 | 0.04 | 0.39 |
| RABC6 | 23 | 234 | 3610 | 18045 | 1797 | 1 | -0.25 | -0.39 | -0.11 |
| RABC7 | 123 | 661 | 3610 | 18045 | 2154 | 0 | 0.61 | 0.52 | 0.70 |
| RABC12 | 28 | 193 | 3523 | 16398 | 1145 | 7.87E-01 | -0.05 | -0.20 | 0.11 |
| RABC14 | 28 | 101 | 3523 | 16398 | 949 | 1 | -0.51 | -0.72 | -0.30 |
| RABC15 | 29 | 573 | 3520 | 20292 | 2047 | 0 | 0.64 | 0.54 | 0.74 |
| RABC16 | 21 | 147 | 3502 | 16077 | 727 | 5.38E-02 | 0.12 | -0.06 | 0.30 |
| RABC18 | 173 | 430 | 3748 | 20694 | 2445 | 7.34E-04 | 0.14 | 0.03 | 0.25 |
| RABC39 | 513 | 522 | 3551 | 17389 | 2094 | 0 | 0.39 | 0.28 | 0.49 |
| RABC68 | 33 | 558 | 3114 | 21500 | 1863 | 0 | 0.86 | 0.76 | 0.96 |
| RABC72 | 30 | 215 | 3518 | 16180 | 1005 | 1.85E-03 | 0.18 | 0.03 | 0.33 |
| RABC73 | 101 | 709 | 3519 | 21792 | 1885 | 0 | 1.00 | 0.90 | 1.09 |
| RABC76 | 133 | 347 | 3683 | 15564 | 1170 | 0 | 0.44 | 0.31 | 0.56 |
| RABC79 | 147 | 903 | 3743 | 31303 | 3504 | 0 | 0.88 | 0.80 | 0.96 |

### Consistent enrichment results are observed by the Meta-analysis

Due to the wide range of variation in the number of background genes and the number of DEGs across datasets, it showed a large degree of heterogeneity with p-value less than 0.05. Therefore, a random effect model was used to pool the effect sizes of all studies. The logarithmic effect point estimate for multiple studies was 0.48 and the 95% confidence interval was [0.17, 0.80]. The combined effect size was significantly higher than that of the randomized case (p = 0.0012). That is to say, consistent enrichment results are observed by the inverse-variance Meta-analysis (Figure 2).

**Figure 2.**
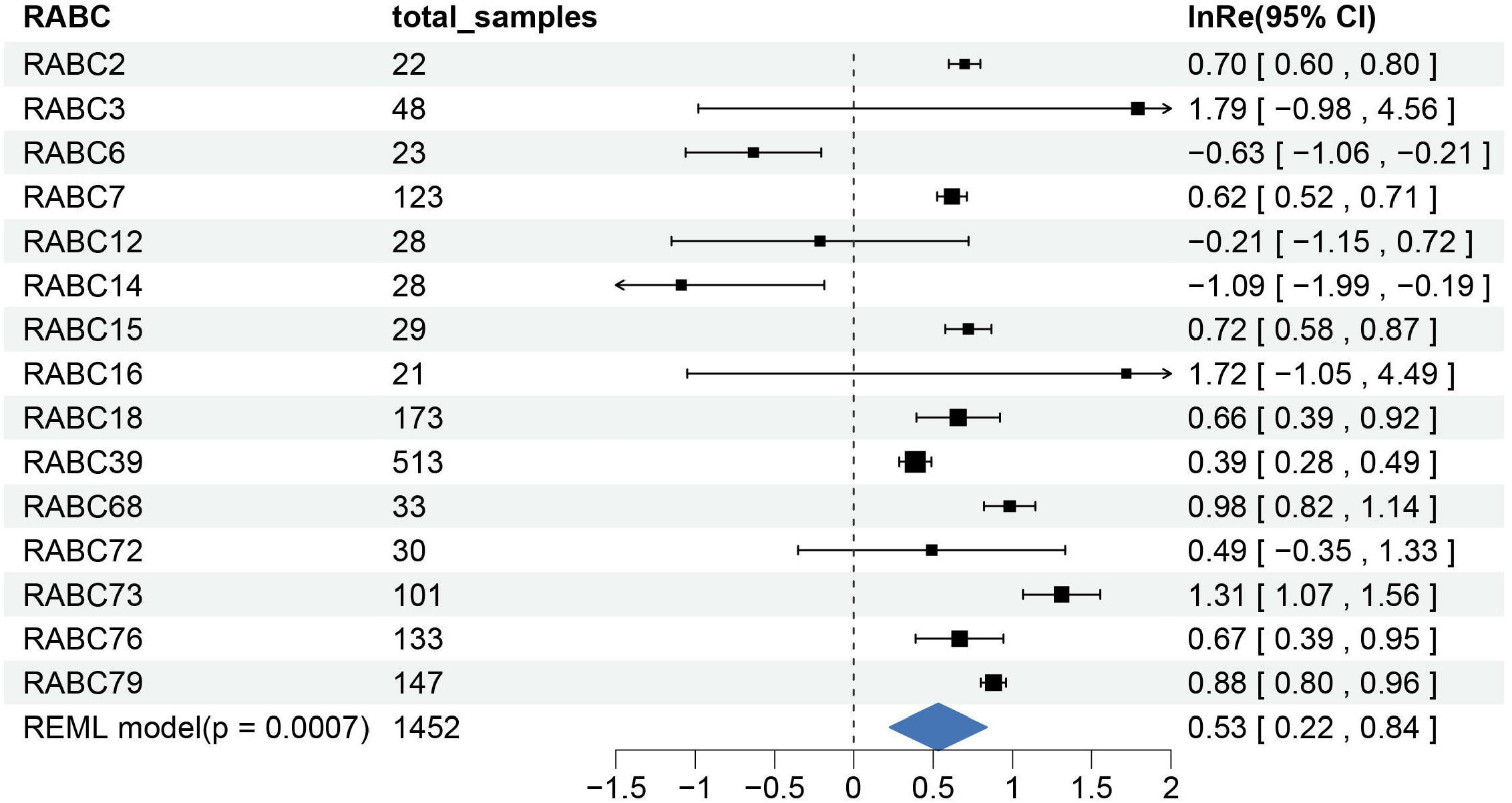
DEGs between RA and controls are significantly enriched to HK genes shown by the forest diagram. The point estimate and 95% confidence interval of effect size for each single dataset was represented by black rectangles and line segment respectively. The point estimate of integrated effect size was represented by a dark blue diamond.

Then we evaluated the publication bias across all these individual studies by funnel diagram, Begg’s and Egger’s tests. Significant publication bias was not observed with p value of Begg’s and Egger’s tests greater than 0.05 (p = 0.59 for Begg’s test and p = 0.48 for Egger’s test).

Besides the stability of the meta-analysis was assessed by the sensitivity analysis. Specifically, each individual study was removed and re-performed the meta-analysis. Low sensitivity was observed by comparing the effect size bias between each new Meta-analysis and the previous overall Meta-analysis (Figure 3). The integration of effect size was still statistically significant with p values less than 0.01 after removing any of the datasets. This further indicates high reliability of consistent enrichment results of Meta analysis.

**Figure 3.**
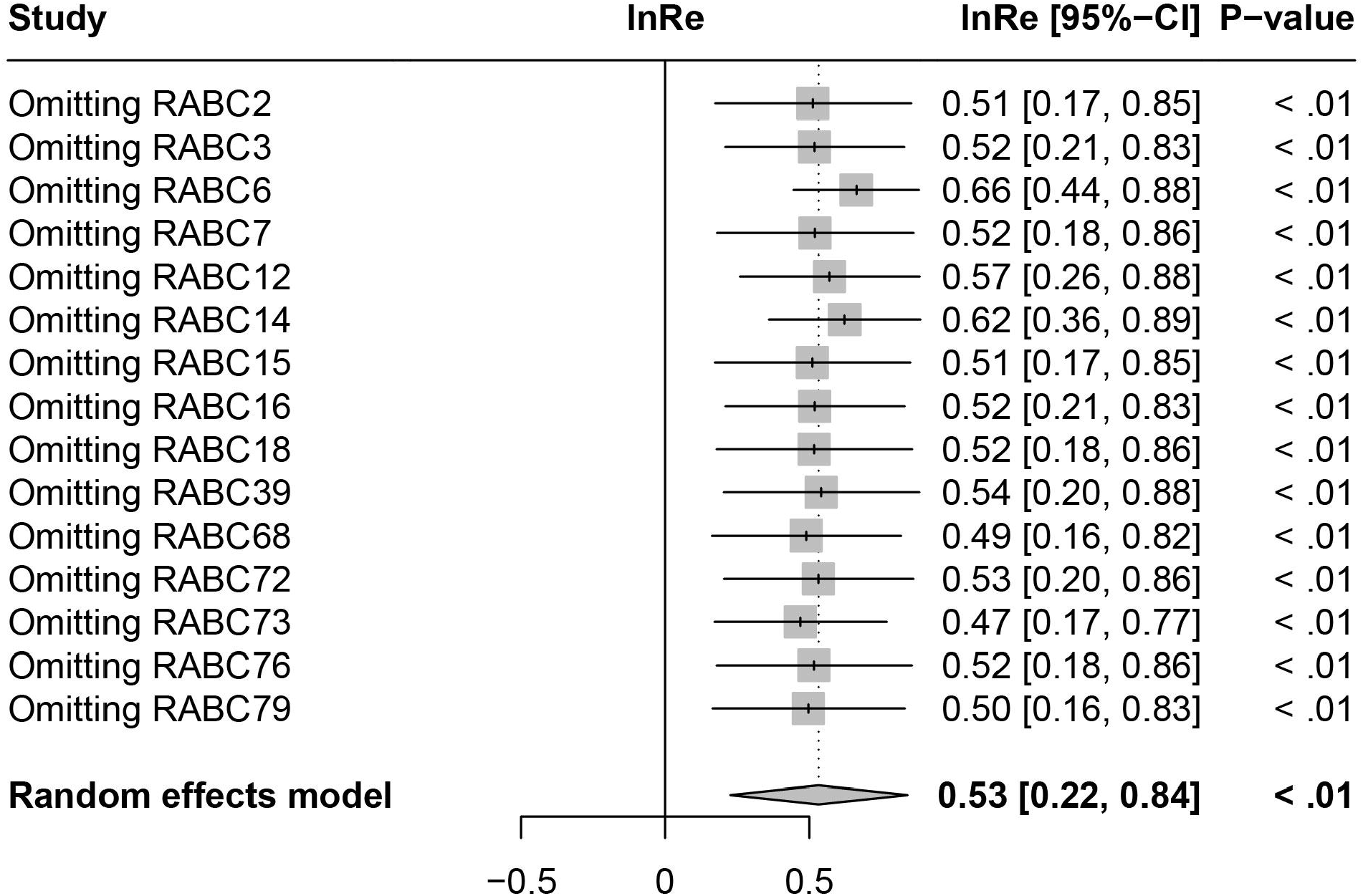
Sensitivity analysis result by removing one dataset each time. A meta-analysis was re-performed by removing each of the datasets and a new estimate and confidence interval of integrated effect size was plotted. The p values of new meta-analyses were placed on the last column.

### HK genes showed higher variance in RA patients compared to healthy controls

For the 15 datasets included in this research, variance, standard deviation (SD) and coefficient of variation (CV) of expression levels of HK genes were calculated for RA samples and healthy controls respectively. The results of two-thirds of the datasets showed that the expression fluctuation of HK genes in RA samples was significantly higher than that of healthy controls with p values of Wilcoxon rank sum tests less than 0.05 (Figure 4, Figure S1-S2). We obtained consistent results for variance, SD and CV indicating that HK genes have aberrant changes in expression levels in RA patients.

**Figure 4.**
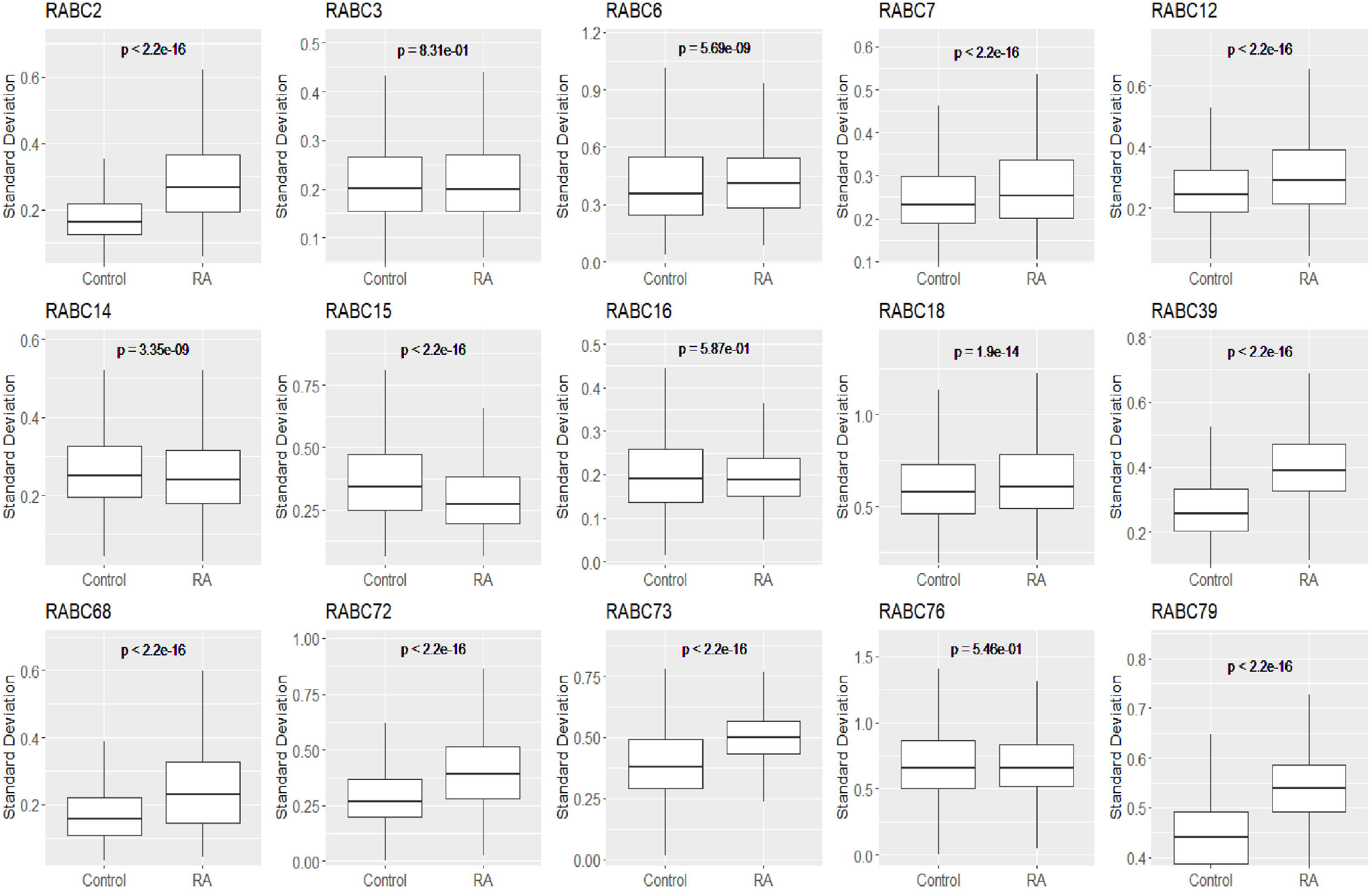
Comparison of the stability of gene expression levels between RA samples and healthy controls. The vertical axis represents the standard deviation of gene expression levels. A P value of Wilcoxon rank sum test is placed on the top of each boxplot.

## Discussion

Housekeeping (HK) genes are generally considered to be a class of genes with stable expression regardless of tissue, cell, or disease state. Therefore HK genes are often used as reference controls to measure the expression levels of other genes. However, through comparative analysis, it was found for the first time that HK genes showed significant expression alterations in RA patients, suggesting that HK genes are involved in the development of RA to a certain extent. This is a great challenge to previous views of housekeeping genes.

It is known that the reliability of data analysis is largely reflected by repeatability. In this study we analyzed multiple datasets of RA gene expression profile using different methods and obtained consistent conclusions that HK genes were significantly enriched in DEGs between RA samples and healthy controls, implying the stability and reliability of the results. Specifically, significant results were observed in 80% (12/15) of the datasets through the cumulative hypergeometric tests. Next the integrated effect size was significantly different compared to that of the randomized situation through a meta-analysis strategy. In other words, the meta-analysis yielded results consistent with the previous cumulative hypergeometric tests, which increased the sample size, reduced the sampling error and further confirmed the reproducibility of the findings.

Considering the possible influence of tissue origin on the results of the analysis, a subgroup analysis was performed in which 14 blood sample datasets were retained while 1 set of synovial tissue samples was removed. The observations of the subgroup analysis were highly consistent with the original results in the manuscript where p value was less than 0.0001, and the point estimation of lnRe is 0.66 with the 95% confidence interval ranging from 0.44 to 0.88 suggesting the reliability and reproducibility of this research (Figure S3).

To further verify the reliability of the results, another standard error estimation method for effect size (relative enrichment, Re) has been applied and the effect size was re-integrated through Meta-analysis (Methods and materials). The combined effect size obtained from the supplementary analysis method was significantly greater than that of the randomized case (p = 0.0015), the point estimate of lnRe was 0.35 and the 95% confidence interval ranged from 0.14 to 0.57 (Figure S4). The hypergeometric distribution background generated by 100,000 random perturbations ensures the accuracy of the standard error estimation, which verifies the stability and reliability of the results from another perspective.

In the present study we obtained a preliminary finding that altered transcript levels of HK genes were observed in RA patients. By calculating and comparing coefficients of variation, standard deviations and variances, we found that in 75% of the datasets, the expression fluctuation of HK genes was significantly greater in RA patients than in healthy controls. Studies have shown that DNA methylation in gene promoter regions plays an important role in the regulation of gene expression and is closely related to RA[23, 24]. Therefore, we wonder whether the relationship between DNA methylation and gene expression of housekeeping genes differs from that of other genes for RA patients. Based on this consideration, we calculated the Pearson correlation coefficients between DNA methylation and gene expression for both housekeeping genes and non-housekeeping genes in RA samples using sample-matched DNA methylation data (GSE138653) and transcriptome data (GSE138746). In order to reduce the bias due to the gene set size difference between housekeeping genes and non-housekeeping, 100 random samplings were conducted, for each sampling an equal number of non-housekeeping genes were drawn from the background genes and then the correlation between the two gene sets was compared. The correlation between DNA methylation and gene expression of housekeeping genes was significantly higher than that of non-housekeeping genes in 94 random samplings with p < 0.05 (Table S2, Figure S5 - S14). Based on the previous transcriptome data analysis and the relationship between DNA methylation and gene expression, we speculate that the dysregulation of housekeeping genes in the pathogenesis of rheumatoid arthritis may involve epigenetic mechanisms. In subsequent studies, we will further verify the observation by large-scale data analysis and experiments so as to elaborate the molecular mechanisms of RA.

### Availability of data and materials

The raw datasets of gene expression profile analyzed during the current study are available in the RABC database [http://www.onethird-lab.com/RABC/].

The housekeeping gene list can be found in the Table S3.

## Ethics approval and consent to participate

Not applicable.

## Consent for publication

Not applicable.

## Author contributions

**WH.L**: Methodology, Validation, Writing – original draft. **H.T**: Methodology, Software, Visualization.**SH.Z**: Methodology, Visualization. **GL.F**: Data analysis, Visualization. **ZW.S**: Data curation. **MM.Z**: Review. **R.C**: Methodology. **XS.C:** Data curation. **X.M** Data curation. **HC. L**: Conceptualization, Supervision, Review and editing. **RJ.Z**: Conceptualization, Project administration, Supervision, Writing – review & editing.

## Acknowledgments

Not applicable.

