## Supplementary figures for "Housekeeping genes are enriched in rheumatoid arthritis-related genes"

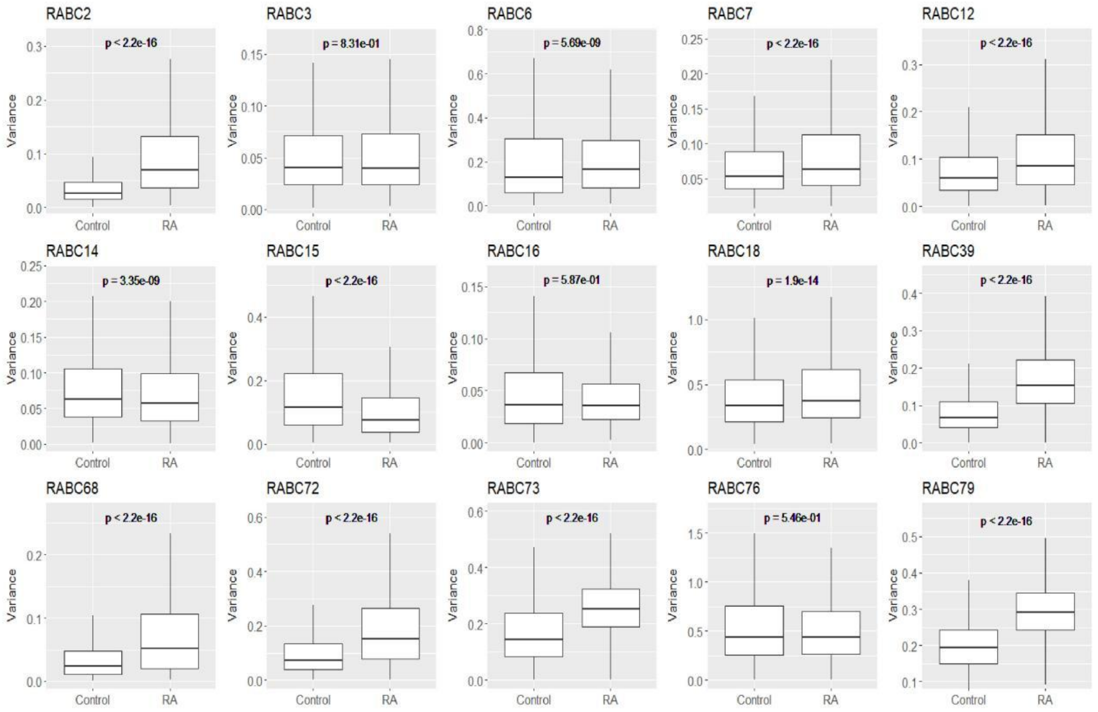


**Figure S1.** Comparison of the stability of gene expression levels between RA samples and healthy controls. The vertical axis represents the variance of gene expression levels. A P value of Wilcoxon rank sum test is placed on the top of each boxplot.

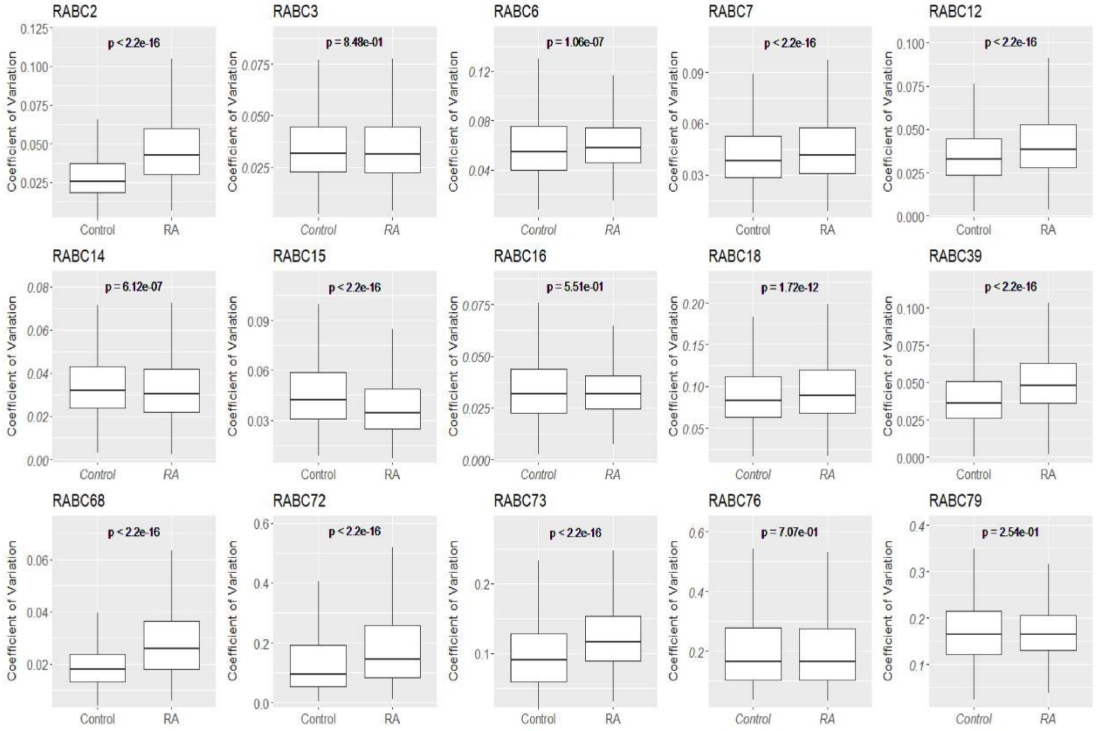


**Figure S2.** Comparison of the stability of gene expression levels between RA samples and healthy controls. The vertical axis represents the coefficient of variation of gene expression levels. A P value of Wilcoxon rank sum test is placed on the top of each boxplot.

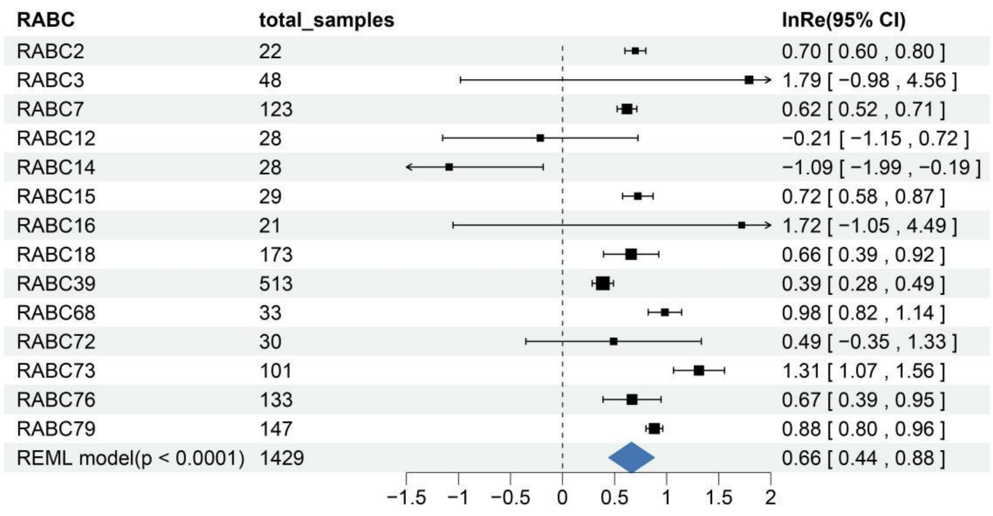


**Figure S3.** Integration of effect size (lnRe) by a subgroup analysis for blood sample datasets. The point estimate and 95% confidence interval of effect size for each single dataset was represented by black rectangles and line segment respectively. The point estimate of integrated effect size was represented by a dark blue diamond.

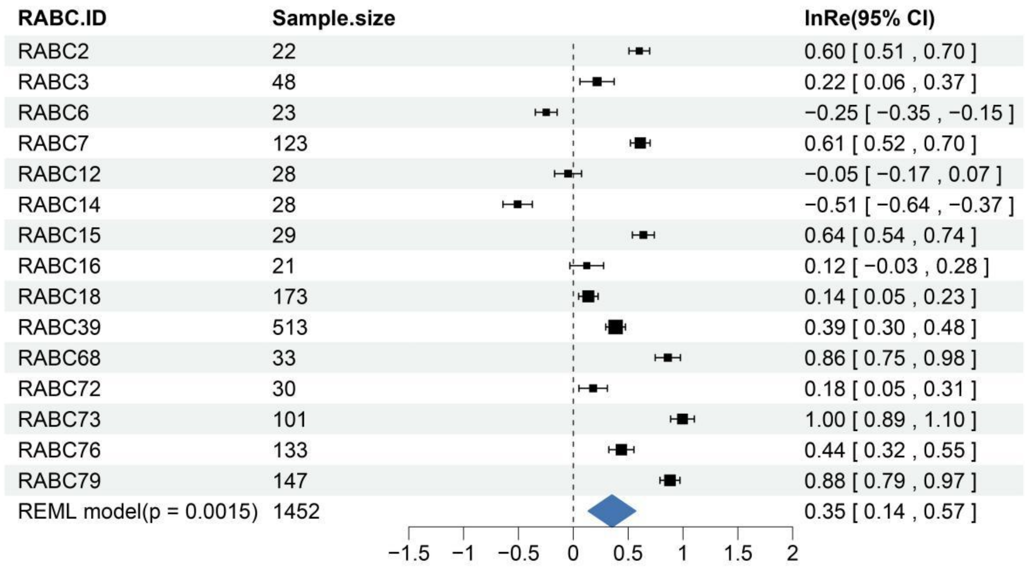


**Figure S4.** Integration of effect size (lnRe) based on the results of another standard error estimation method. The point estimate and 95% confidence interval of effect size for each single dataset was represented by black rectangles and line segment respectively. The point estimate of integrated effect size was represented by a dark blue diamond.

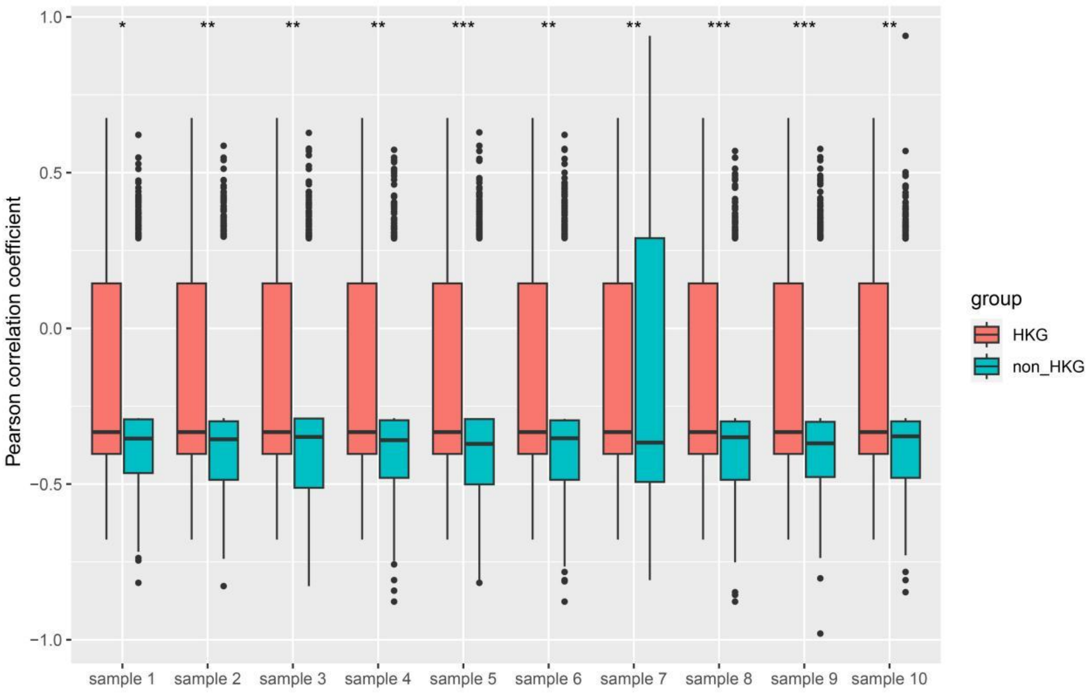


**Figure S5.** Housekeeping genes have higher Pearson correlation coefficient than non-housekeeping genes in RA patients (Random sampling from the 1st to the 10th). One asterisk (*) means that the p-value is less than 0.05, and two asterisks (**), three asterisks (***), and four asterisks (****) indicate that the p-value is less than 0.01, 0.001, and 0.0001, respectively. The sign of 'ns' means 'not significant'.

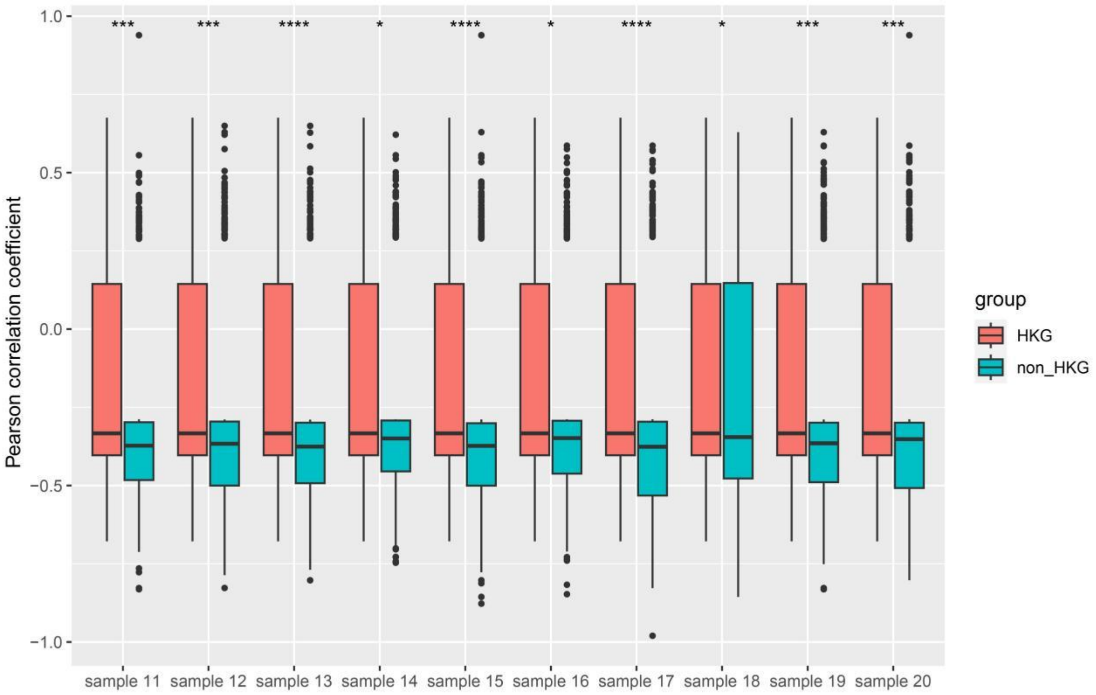


**Figure S6.** Housekeeping genes have higher Pearson correlation coefficient than non-housekeeping genes in RA patients (Random sampling from the 11th to the 20th). One asterisk (*) means that the p-value is less than 0.05, and two asterisks (**), three asterisks (***), and four asterisks (****) indicate that the p-value is less than 0.01, 0.001, and 0.0001, respectively. The sign of 'ns' means 'not significant'.

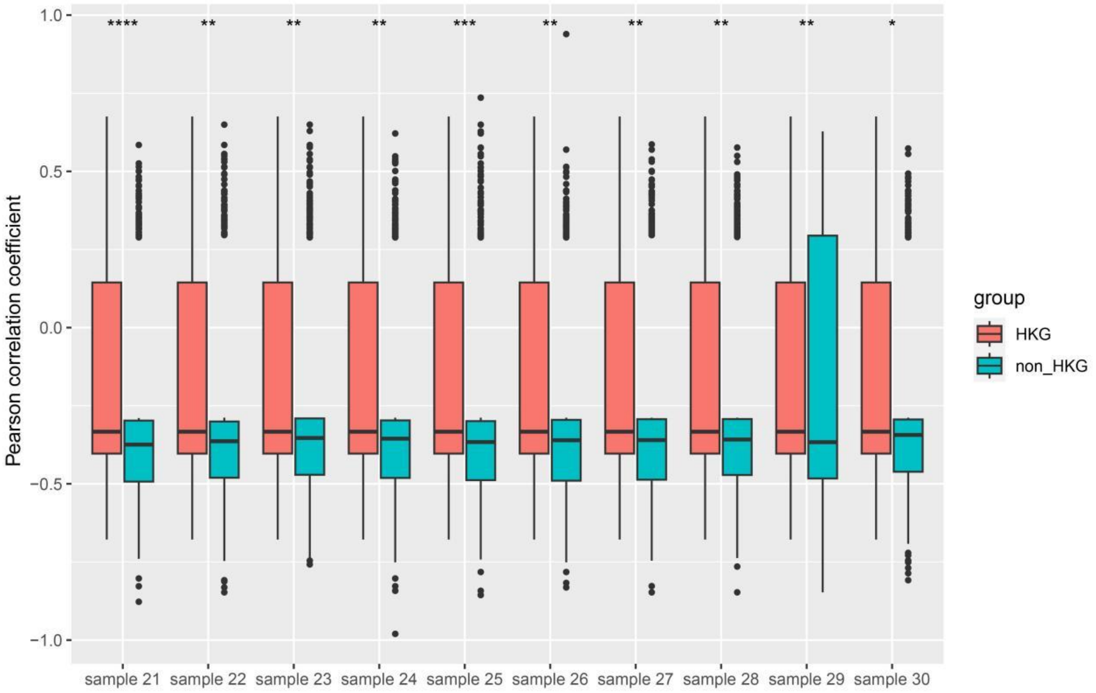


**Figure S7.** Housekeeping genes have higher Pearson correlation coefficient than non-housekeeping genes in RA patients (Random sampling from the 21st to the 30th). One asterisk (*) means that the p-value is less than 0.05, and two asterisks (**), three asterisks (***), and four asterisks (****) indicate that the p-value is less than 0.01, 0.001, and 0.0001, respectively. The sign of 'ns' means 'not significant'.

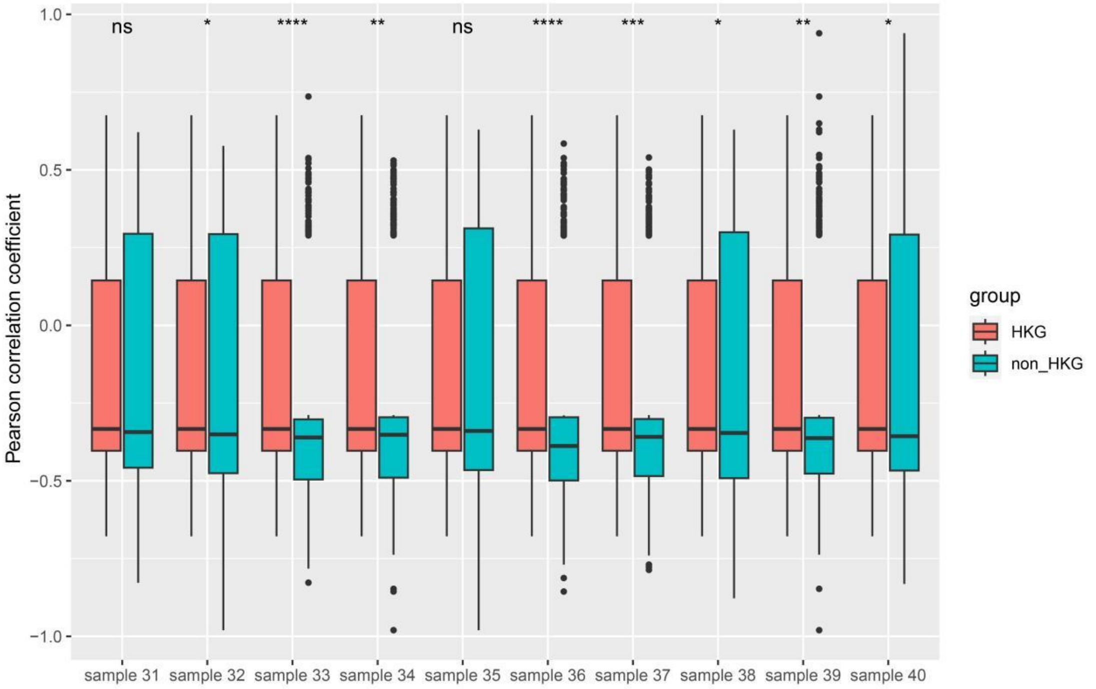


**Figure S8.** Housekeeping genes have higher Pearson correlation coefficient than non-housekeeping genes in RA patients (Random sampling from the 31st to the 40th). One asterisk (*) means that the p-value is less than 0.05, and two asterisks (**), three asterisks (***), and four asterisks (****) indicate that the p-value is less than 0.01, 0.001, and 0.0001, respectively. The sign of 'ns' means 'not significant'.

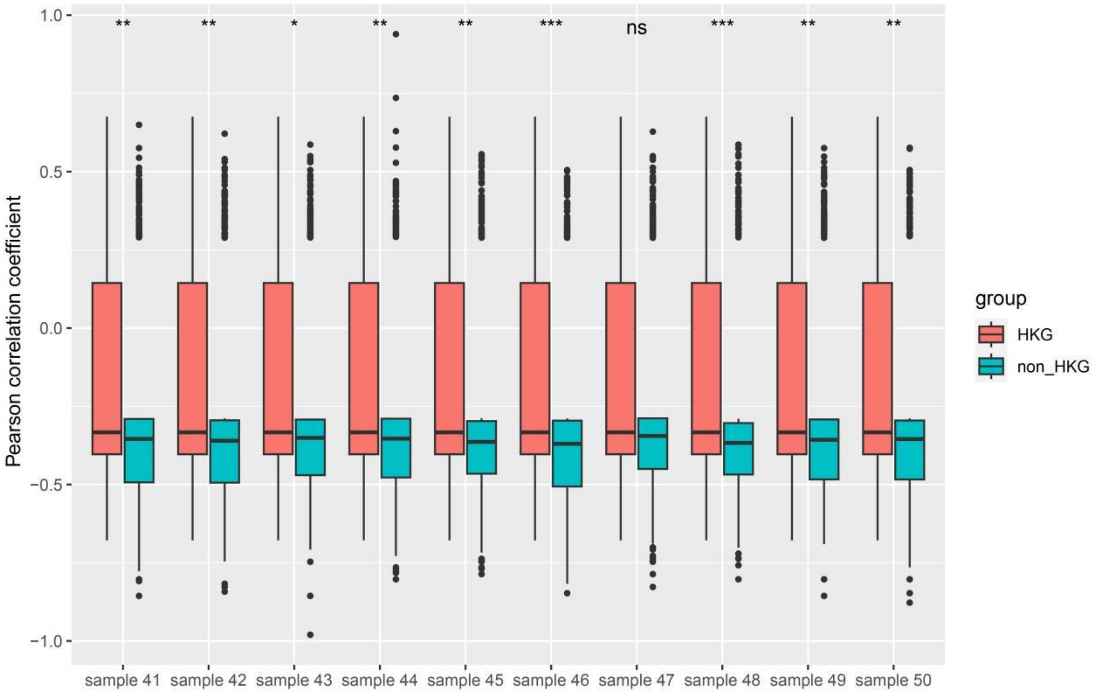


**Figure S9.** Housekeeping genes have higher Pearson correlation coefficient than non-housekeeping genes in RA patients (Random sampling from the 41st to the 50th). One asterisk (*) means that the p-value is less than 0.05, and two asterisks (**), three asterisks (***), and four asterisks (****) indicate that the p-value is less than 0.01, 0.001, and 0.0001, respectively. The sign of 'ns' means 'not significant'.

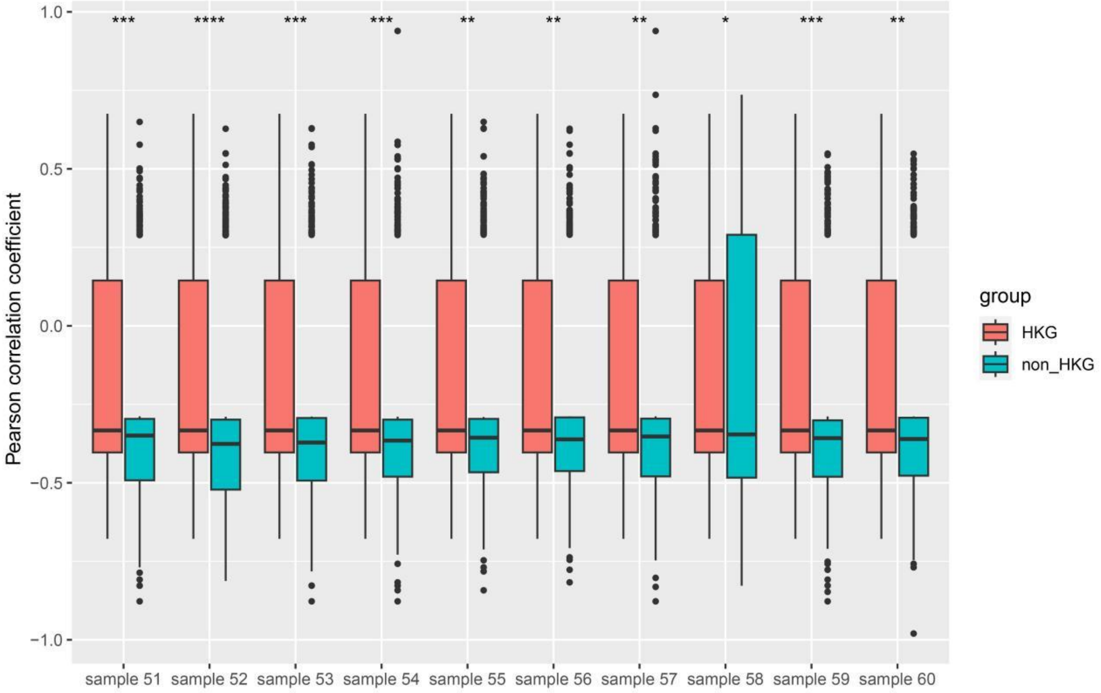


**Figure S10.** Housekeeping genes have higher Pearson correlation coefficient than non-housekeeping genes in RA patients (Random sampling from the 51st to the 60th). One asterisk (*) means that the p-value is less than 0.05, and two asterisks (**), three asterisks (***), and four asterisks (****) indicate that the p-value is less than 0.01, 0.001, and 0.0001, respectively. The sign of 'ns' means 'not significant'.

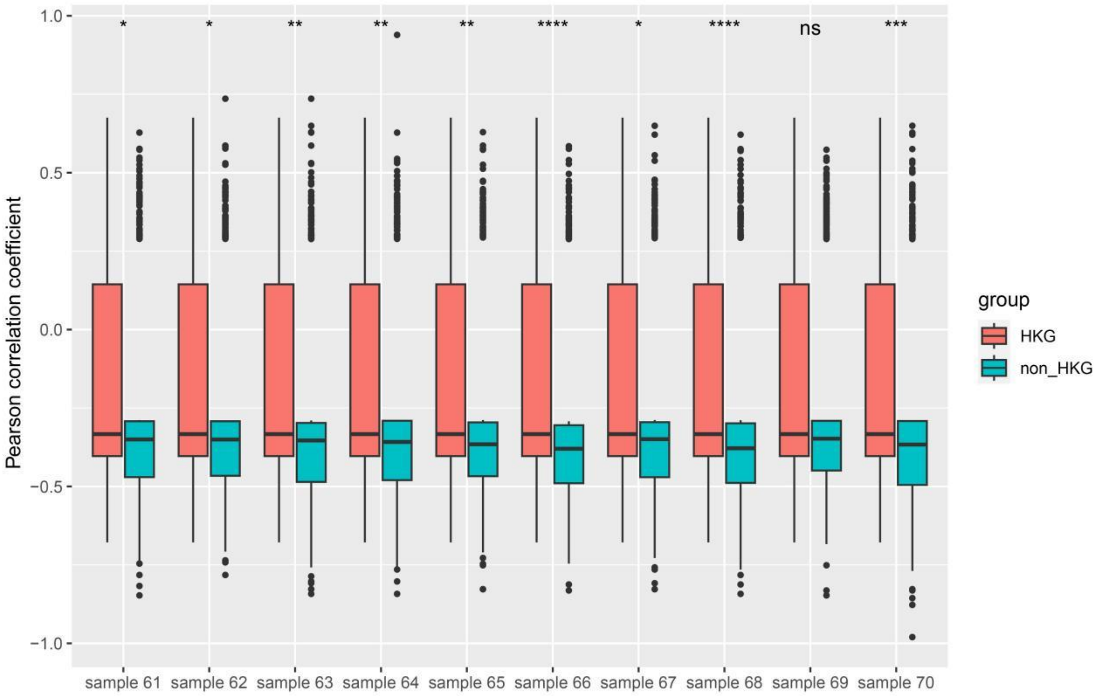


**Figure S11.** Housekeeping genes have higher Pearson correlation coefficient than non-housekeeping genes in RA patients (Random sampling from the 61st to the 70th). One asterisk (*) means that the p-value is less than 0.05, and two asterisks (**), three asterisks (***), and four asterisks (****) indicate that the p-value is less than 0.01, 0.001, and 0.0001, respectively. The sign of 'ns' means 'not significant'.

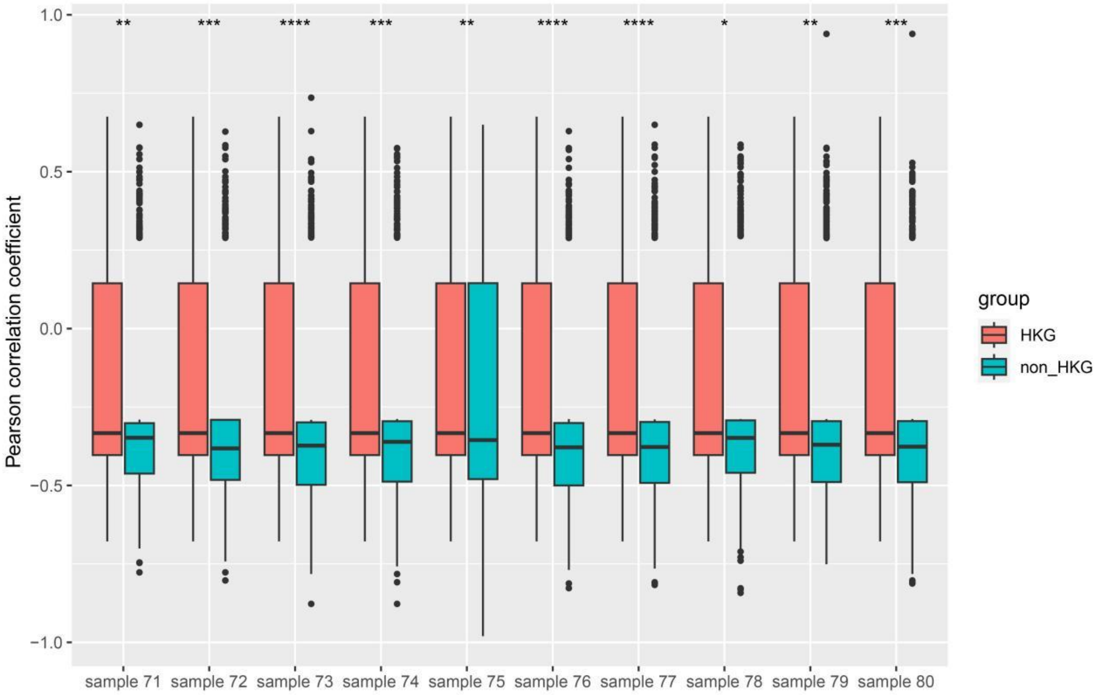


**Figure S12.** Housekeeping genes have higher Pearson correlation coefficient than non-housekeeping genes in RA patients (Random sampling from the 71st to the 80th). One asterisk (*) means that the p-value is less than 0.05, and two asterisks (**), three asterisks (***), and four asterisks (****) indicate that the p-value is less than 0.01, 0.001, and 0.0001, respectively. The sign of 'ns' means 'not significant'.

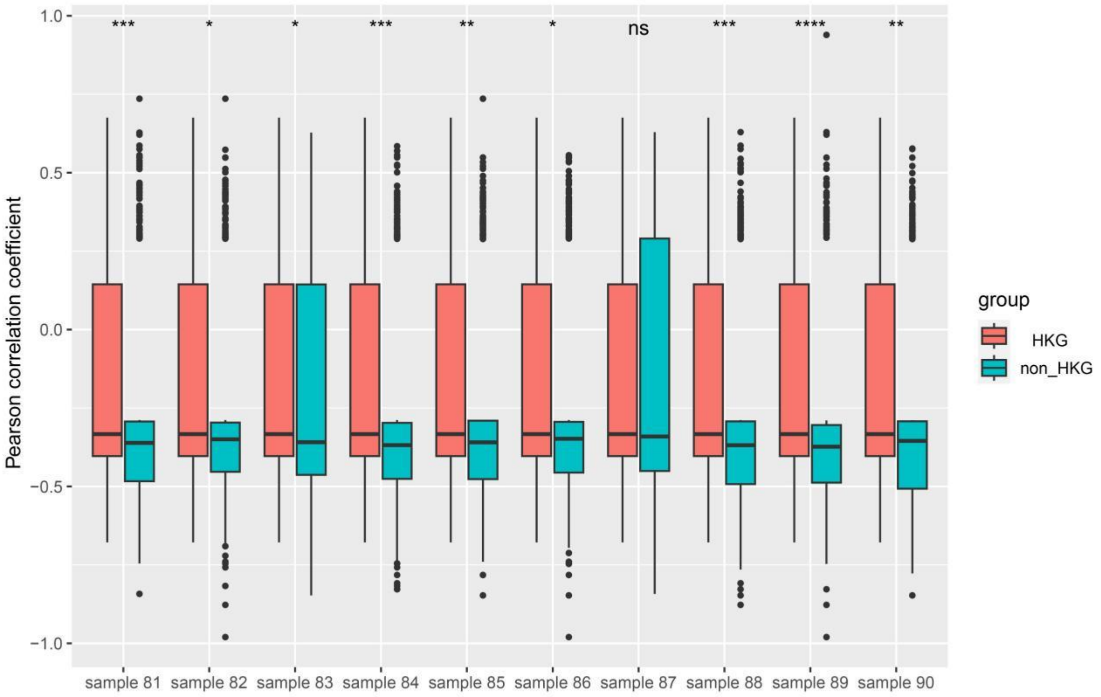


**Figure S13.** Housekeeping genes have higher Pearson correlation coefficient than non-housekeeping genes in RA patients (Random sampling from the 81st to the 90th). One asterisk (*) means that the p-value is less than 0.05, and two asterisks (**), three asterisks (***), and four asterisks (****) indicate that the p-value is less than 0.01, 0.001, and 0.0001, respectively. The sign of 'ns' means 'not significant'.

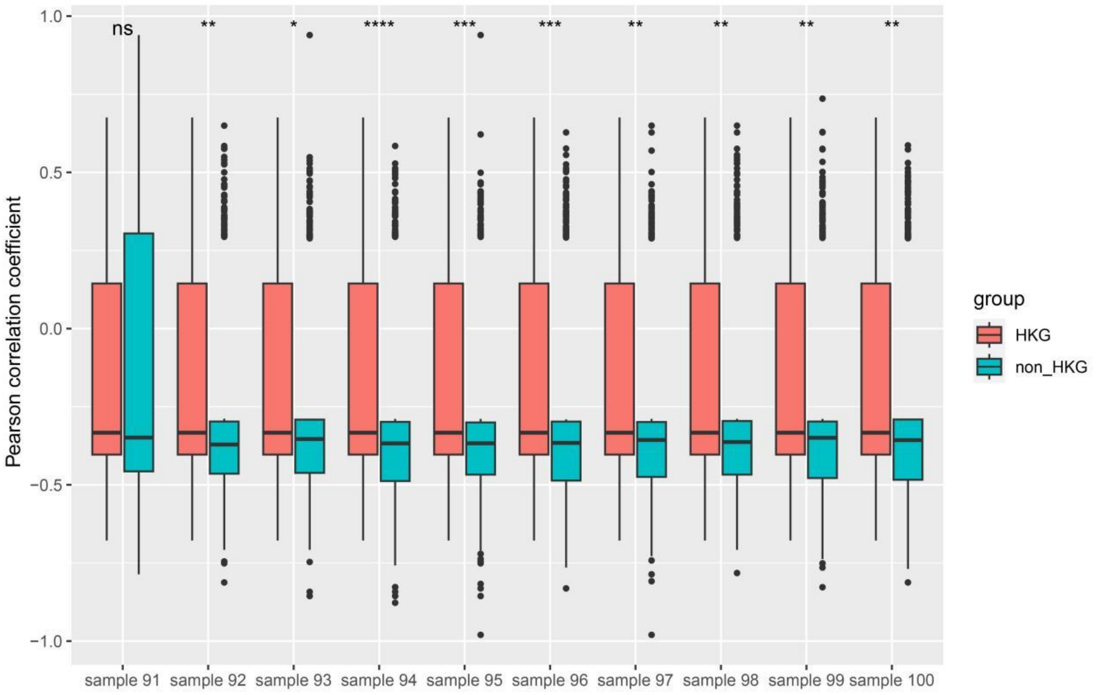


**Figure S14.** Housekeeping genes have higher Pearson correlation coefficient than non-housekeeping genes in RA patients (Random sampling from the 91st to the 100th). One asterisk (*) means that the p-value is less than 0.05, and two asterisks (**), three asterisks (***), and four asterisks (****) indicate that the p-value is less than 0.01, 0.001, and 0.0001, respectively. The sign of 'ns' means 'not significant'.
